# Stochastic microbial community assembly decreases biogeochemical function

**DOI:** 10.1101/183897

**Authors:** Emily B. Graham, James C. Stegen

## Abstract

Ecological mechanisms influence relationships among microbial communities, which in turn impact biogeochemistry. In particular, microbial communities are assembled by deterministic (e.g., selection) and stochastic (e.g., dispersal) processes, and the relative influence of these two process types is hypothesized to alter the influence of microbial communities over biogeochemical function, which we define generically to represent any biogeochemical reaction of interest. We used an ecological simulation model to evaluate this hypothesis. We assembled receiving communities under different levels of dispersal from a source community that was assembled purely by deterministic selection. The dispersal scenarios ranged from no dispersal (i.e., selection-only) to dispersal rates high enough to overwhelm selection (i.e., homogenizing dispersal). We used an aggregate measure of community fitness to infer its biogeochemical function relative to other communities. We also used ecological null models to further link the relative influence of deterministic assembly to function. We found that increasing rates of dispersal decrease biogeochemical function by increasing the proportion of maladapted taxa in a local community. Niche breadth was also a key determinant of biogeochemical function, suggesting a tradeoff between the function of generalist and specialist species. Together, our results highlight the influence of spatial processes on biogeochemical function and indicate the need to account for such effects in models that aim to predict biogeochemical function under future environmental scenarios.

## 1. Introduction

Recent attempts to link microbial communities and environmental biogeochemistry have yielded mixed results [1–6], leading researchers to propose the inclusion of community assembly mechanisms such as dispersal and selection in our understanding of biogeochemistry [2,7–9]. Although much work has examined how assembly processes influence the maintenance of diversity and other ecosystem-level processes in macrobial systems [10–13], our comprehension of how these processes influence microbially-mediated biogeochemical cycles is still nascent [2,8,14]. Thus, there is a need to discern the circumstances under which knowledge on assembly processes is valuable for predicting biogeochemical function.

Community assembly processes collectively operate through space and time to determine microbial community composition [3,7,14,15]. They fall into two predominate categories that can be summarized as influenced (i.e., deterministic) or uninfluenced (i.e., stochastic) by abiotic and abiotic environmental conditions. Stochastic processes can be further classified into dispersal, evolutionary diversification, and ecological drift, while determinism is largely dictated by selection [7,16]. Experimental research has shown unpredictable relationships between microbial diversity and biogeochemical function (generically defined here to represent any biogeochemical reaction of interest), leading to the hypothesis that differences in community assembly history—and thus the relative contributions of stochastic and deterministic processes—drives relationships between microbial community structure and biogeochemical function [8,9].

Dispersal in particular may vary the relationship between community structure and biogeochemical function [7]. Both positive and negative relationships have been hypothesized [reviewed in 17]. The ‘portfolio effect’ argues for enhanced community functioning under high levels of dispersal, proposing that high diversity communities are more likely to contain more beneficial species properties on average than lower diversity communities [18,19]. Additionally, if dispersal increases biodiversity, there should be a greater chance that the community can occupy more niches (*i.e.*, niche complementarity), reducing direct competition and increasing function [20].

Alternatively, dispersal may decrease community-level biogeochemical function [7,21]. High rates of dispersal can add organisms to a microbial community that are not well-suited to local environmental conditions [i.e., mass effect or source-sink dynamics 22,23]. Maladapted individuals may invest more in cell maintenance to survive as opposed to investing in cellular machinery associated with biogeochemical reactions needed to obtain energy for growth and reproduction. In this case, the community’s ability to drive biogeochemical reactions may be depressed. For instance, pH [24] and salinity [25,26] are widely considered strong regulators of microbial community structure. If microorganisms are well adapted to and disperse from a moderate pH or salinity environment to a more extreme environment, they may be maladapted and have to expend energy to express traits that maintain neutral internal pH (e.g., H+ pumps) or maintain cellular water content (e.g., osmotic stress factors). These cell maintenance activities detract from the energy available to transform biogeochemical constituents and may suppress overall community rates of biogeochemical function. In contrast, locally adapted species would putatively have more efficient mechanisms for cell maintenance in the local environment and be able to allocate more energy for catalyzing biogeochemical reactions.

These dispersal effects also interact with local selective pressures and the physiological ability of organisms to function across a range of environments to collectively influence biogeochemical function in uncertain ways. Here, we propose that (1) more stochastically assembled communities are composed of species that are less well adapted to the local environment and, in turn, that (2) stochastic assembly processes decrease biogeochemical function (Figure 1). Our aim is to formalize these hypotheses and provide a simulation-based demonstration of how stochastic assembly can influence function. To do so, we employ an ecological simulation model to explicitly represent dispersal and selection-based processes, and we leverage ecological null models that have a long history of use in inferring assembly processes [27]. We link the resulting communities to biogeochemical function through organismal fitness. In our conceptualization, biogeochemical function is a generic representation, and thus, our results can be applied to any process of interest.

**Figure 1.**
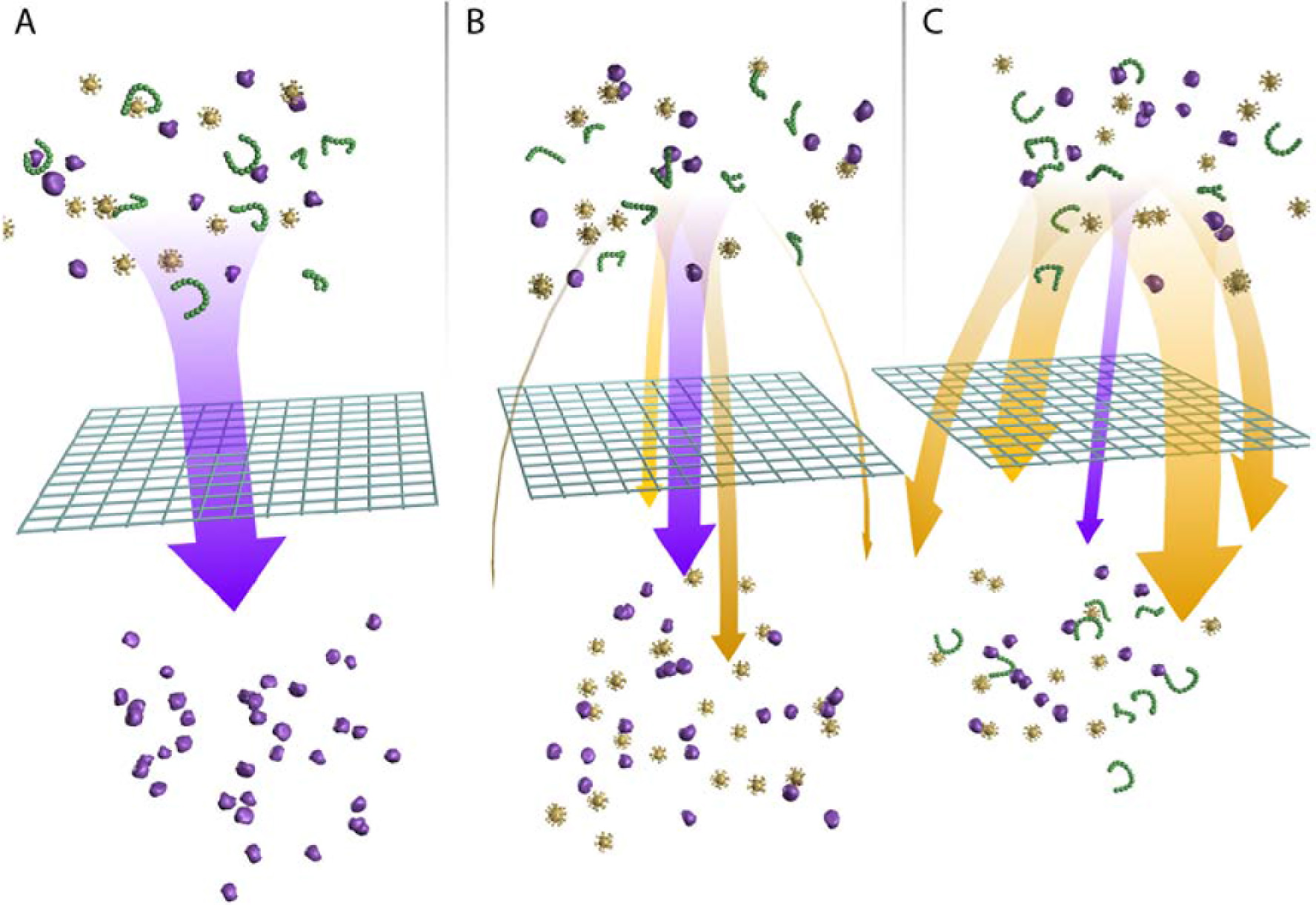
We propose a conceptual model in which stochastic assembly processes decrease biogeochemical function. Purple organisms in this figure represent all species that are well-adapted to and are thus good competitors in a given environment. Yellow and green organisms represent all species that that are less adapted to the environment than purple organisms. While not displayed for simplicity, we conceptualize multiple species within each color. We acknowledge that the environment influences microbial community composition through effects of both abiotic (e.g., resource availability) and biotic (e.g., competition and predator-prey interactions) factors. We use the term ‘selective filter’ to indicate influences of both factors on an organism’s fitness [28]. **(A)** In a community structured entirely by determinism, selective filtering restricts community composition to species that are well-adapted to prevailing conditions, resulting in enhanced biogeochemical function. **(B)** In communities with moderate stochasticity (here, moderate rates of dispersal), there is an increase in the abundance of maladapted organisms in the community. In turn, the community is less efficient and exhibits lower biogeochemical function. **(C)** Under high levels of stochasticity (here, high rates of dispersal), a large portion of community members are maladapted, resulting in the lowest rates of biogeochemical function.

## 2. Materials and Methods

All simulations, null models, statistical analyses, and graphics were completed in *R* software (https://cran.r-project.org/). The simulation model consisted of two parts and were followed by statistical analysis. The model builds upon previous work by Stegen, Hulbert, Graham, and others [2,14,15,29–32]. Relative to this previous work, the model used here is unique in connecting evolutionary diversification, variation in the relative influences of stochastic and deterministic processes, null models to infer those influences, and biogeochemical function. Previous models have addressed some subset of those features (e.g., connecting evolutionary processes with stochastic and deterministic ecology), but as far as we are aware, previous studies have not integrated all features examined here. One hundred replicates were run for each parameter combination in the simulation model. A central purpose of the simulation model was to vary the influences of community assembly processes. Previously developed null models (see below) were used to identify parameter combinations that provided a range of scenarios across which the relative balance among community assembly processes varied. As such, parameter values were selected to generate conceptual outcomes needed to evaluate the relationship between assembly processes and biogeochemical function. Specific parameter values do not, therefore, reflect conditions in any particular ecosystem.

### 2.1 Regional Species Pool Simulation

First, a regional species was constructed following the protocol outlined in Stegen *et al.* [15]. Regional species pools were constructed by simulating diversification in which entirely new species arise through mutations in the environmental optima of ancestral organisms. Environmental optima evolve along an arbitrary continuum from 0 to 1, following a Brownian process. Regional species pools reach equilibria according to the constraints described by Stegen *et al.* [15] and Hurlbert and Stegen [29] and summarized here: (1) we define a maximum number of total individuals in the pool (2 million) such that the population size of a given species declines with an increasing number of species, and (2) the probability of extinction for a given species increases as its population size decreases according to a negative exponential function [population extinction probability ∝ exp(−0.001 *population size)].

The evolution of a regional species pool was initiated from a single ancestor with a randomly chosen environmental optimum (initially comprising all 2 million individuals in the population). Mutation probability was set as 1.00E-05. A descendant’s environmental optimum deviated from its ancestor by a quantity selected from a Gaussian distribution with mean 0 and standard deviation 0.2. Following mutation, population sizes were reduced evenly such that the total number of individuals remained 2 million. The simulation was run for 250 time steps, which was sufficient to reach equilibrium species richness.

### 2.2 Community Assembly

The model’s second component assembled 4 local communities from the regional species pool according to scenarios conceptualized to test our hypotheses. Species were drawn from the regional species pool to generate a source community under weak selection and three receiving communities with no dispersal, moderate dispersal, and high dispersal in which organismal niche breadth (*n*, 0.0075 to 0.175) and environmental conditions (*E*, 0.05 to 0.95) were allowed to vary across simulations. A simplifying assumption of the model was that all organisms in a simulation had equivalent niche breadth. The purpose of this assumption was to examine tradeoffs between communities comprised of high-functioning, specialist organisms vs. those comprised of lower-functioning, generalist species. Our intent was to simulate communities across a gradient in the degree of specialization (i.e., niche breadth). This allowed for an evaluation of the influence of niche breadth on the relationship between assembly processes and biogeochemical function. All communities had 100 species and 10,000 individuals, drawn probabilistically from the regional species pool. To define species presence/absence in each community we drew 100 species without replacement from the regional species pool based on selection probabilities described below. In turn, we drew 10,000 individuals with replacement into those 100 species using the same selection probabilities. Selection probabilities (*P*) of each species from the regional pool were set by a Gaussian function with variance *n* (reflecting niche breadth) and the deviation (*d*) of each species environmental optimum from the local environment per the following equation:

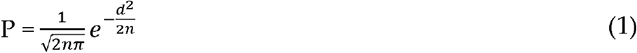

This equation represents the probability of an individual from a given species surviving in a given environment—and thus the strength of selection for or against it—as directly related to three factors: (1) its own environmental optimum, (2) the simulated environment in which it finds itself, and (3) its niche breadth.

For assembly of the source community, we used one niche breadth (*n*) for all simulations, which was the maximum value used for receiving communities (0.175). This value represents generalist organisms, which allows for assembly of species representing a broader range of environmental optima than when niches are narrow. The environmental conditions in the source community was also set to a single value using the following procedure: We generated 10 regional species pools and combined species abundances and environmental optima from these pools to generate one aggregate pool representative of the probable distributions of environmental optima yielded by our simulations. We set the environmental optimum of the source community to one end of this spectrum (5^th^ percentile) to allow for comparisons with receiving communities that had the same or larger environmental values. This allowed us to study emergent behavior across a broad range of environmental differences between the source and receiving communities.

For receiving communities, we allowed the environmental conditions and niche breadth to vary across simulations. Environmental conditions ranged from 0.05 to 0.95 by intervals of 0.04736842 to yield 20 conditions. Niche breadth ranged from 0.0075 to 0.175 by 0.008815789 to yield 20 conditions. Receiving communities were assembled under all possible combinations of environmental conditions and niche breadths. Communities for the selection-only (i.e., no dispersal from the source community) case were assembled based only on the selection probabilities as defined by Equation 1, using the same approach as for the assembly of the source community. For moderate and homogenizing dispersal, we modified selection probabilities to incorporate species dispersing from the source community as defined by the following equation:

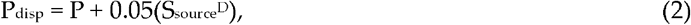

where P_disp_ is the selection probability of a given species accounting for dispersal, S_source_ is the abundance of that species in the source community, and *D* a parameter reflecting dispersal rate. This equation alters the selection probability without dispersal (Eq. 1) with an exponential modifier that enhances the probability of selection for species that are abundant in the source community. Parameter *D* was set to 1 for moderate dispersal and 2 for homogenizing dispersal. All possible communities were simulated with 100 replicate regional species pools such that all possible combination of parameters were used once with each regional species pool.

Our estimation of biogeochemical function is meant to be illustrative and is not associated with any specific reaction. Given this perspective, we make a simplifying conceptual assumption that individuals well-fit to their environment generate higher rates of biogeochemical function than maladapted individuals. The motivation for this assumption is that individuals that are maladapted to a given environmental condition will have to use a larger portion of available energy to maintain their physiological state than well-adapted organisms. In turn, maladapted organisms can invest less in the production of enzymes needed to carry out biogeochemical reactions, thereby leading to lower biogeochemical rates.

In our model, selection probability of a given species in a given environment (Eq. 1) defines how adapted an individual of that species is to its local environment. This leads to another simplifying assumption: the contribution of an individual to the overall biogeochemical rate (*B*) is directly proportional to how well adapted it is to the local environment such that the contribution of each individual is a linear function of its selection probability within a given environment. The biogeochemical contribution of each species is therefore found by multiplying its selection probability by its abundance. To find the total biogeochemical rate for each community we then summed across all species in a community. Biogeochemical function for each community was thus calculated as:

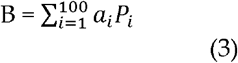
where *B* is the biogeochemical function for a given community and *a*_*i*_ and *P*_*i*_ are the abundance and probability of selection for species *i,* respectively (note there were 100 species within each community). An inherent result of this calculation is that simulations with smaller niche breadth have higher maximum selection probabilities (see Eq. 1), which can lead to higher biogeochemical function, relative to simulations with larger niche breadth. Our formulation therefore assumes higher biogeochemical function for specialist organisms, but only if they are well adapted to their local environment. This assumption reflects a tradeoff between the breadth of environments an individual can persist in and the maximum fitness of an individual in any one environment [discussed in 33].

### 2.3 Ecological Inferences Using Null Models

Following the assembly of communities, the relative influences of stochastic and deterministic assembly processes in structuring communities were estimated using a null modeling approach previous described in Stegen *et al.* [15,30]. We refer the reader to these earlier publications for full details, and provide only a summary of the major elements of the null modeling approach here. The composition of each receiving community was compared to an associated source community that was assembled from the same regional species pool. We first estimated pairwise phylogenetic turnover between a given pair of communities. This was done by calculating the abundance-weighted β-mean-nearest-taxon-distance (βMNTD) [34,35]. A null model was then run 999 times. In each iteration of the null model, species names were moved randomly across the tips of the regional pool phylogeny. This breaks phylogenetic relationships among taxa observed in each community. Using the resulting (randomized) phylogenetic relationships we re-calculated phylogenetic turnover between the pair of communities and refer to this as βMNTD_null_. Running the null model 999 times generated a distribution of βMNTD_null_ values. We then compared the observed (βMNTD to the mean of the (βMNTD_null_ distribution and normalized this difference by the standard deviation of the (βMNTD_null_ distribution. The difference between βMNTD and the (βMNTD_null_ distribution was therefore measured in units of standard deviations and is referred to as the β-nearest taxon index (βNTI) [31]. Values of (βNTI that are < −2 or > +2 are deemed significant in the sense that observed βMNTD deviated significantly from the βMNTD_null_ distribution. The βMNTD_null_ distribution is what’s expected when community assembly is not strongly influenced by deterministic ecological selection. Significant deviation from this distribution therefore indicates that selective pressures are very similar (βNTI < −2) or very different (βNTI > +2) between the two communities being compared. Following the convention of Dini-Andreote et al. [36] we refer to βNTI < −2 as indicating homogeneous selection (i.e., significantly less turnover than expected due to consistent selective pressures) and (βNTI > +2 as indicating variable selection (i.e., significantly more turnover than expected due to divergent selective pressures). Inferences from βNTI have previously been shown to be robust [15]. This method has also been used extensively across a broad range of systems [e.g., 2,14,37–39] and is described in detail in previous work [31].

### 2.4 Statistical Analysis

We analyzed differences in model outputs using standard statistical approaches. We calculated the alpha diversity of each source and receiving community using the Inverse Simpson Index [40] in the *R* package ‘vegan’ [41]. Differences in alpha diversity across communities were evaluated with one-way ANOVA followed by post-hoc Tukey’s HSD tests. We used pairwise Kolmogorov-Smirnov tests to compare distributions of species optima between simulations (all distributions were non-normal). To compare biogeochemical function of the three dispersal cases, we used one-way ANOVA followed by post-hoc Tukey’s HSD tests. We also analyzed how biogeochemical function changed as the environmental difference between source and receiving communities increased; this was done using quadratic regressions due to non-linearity in the relationships. We also compared the influence of dispersal on biogeochemical function across different niche breadths. This was done by first finding the ratio of function in selection-only communities to function in associated homogenizing dispersal communities. Ratios were calculated by comparing communities assembled from the same regional species pool and with identical environmental condition and niche breadth. The resulting distributions of ratios were compared across different niche breadths using one-way ANOVA followed by post-hoc Tukey’s HSD tests. To evaluate the relationship between the relative influence of stochastic assembly (inferred from the value of βNTI) and biogeochemical function, correlations between βNTI and biogeochemical function were assessed with linear regression. In most studies βNTI values are not independent of each other such that statistical significance requires a permutation-based method such as a Mantel test. Here, each βNTI estimate is independent whereby standard statistical methods that assume independence are appropriate.

## 3. Results and Discussion

As ecosystem process models become more sophisticated [e.g., 42,43,44], there is a need to improve these models by better understanding the linkages among community assembly processes and ecosystem function. Here, we used an ecological simulation model to highlight the importance of stochastic microbial community assembly for biogeochemical function. Our results suggest that incorporating assembly processes into ecosystem models may improve model predictions of biogeochemical function under future environmental conditions.

### 3.1 Microbial Community Composition in Response to Stochasticity

We found that diversity was highest when both stochastic and deterministic assembly processes influenced community structure (Figure 2). In communities assembled with moderate to broad niches, intermediate amounts of dispersal led to the highest diversity (Figure 2A, 2B). These moderate-dispersal communities were characterized by distributions of environmental optima (across species and individuals) that did not match the source or selection-only distributions, and instead reflect an influence of both dispersal from the source and local selection (Figure 3, right column). Both moderate- and homogenizing-dispersal cases exhibited higher diversity than source or selection-only communities (Figure 2A, 2B). We note that the slight differences in diversity between source and selection-only communities were due to environmental conditions in source communities being defined at one end of the environmental spectrum. This edge-effect truncated its distribution of species environmental optima, causing the distribution to be right skewed (Figure 3, left column). Our results suggest a conceptual parallel to Connell’s [45] Intermediate Disturbance Hypothesis, whereby intermediate levels of dispersal lead to the highest overall diversity, but only when niche breadth is broad enough to allow for strong contributions from both stochastic and deterministic assembly processes (Figure 2).

**Figure 2.**
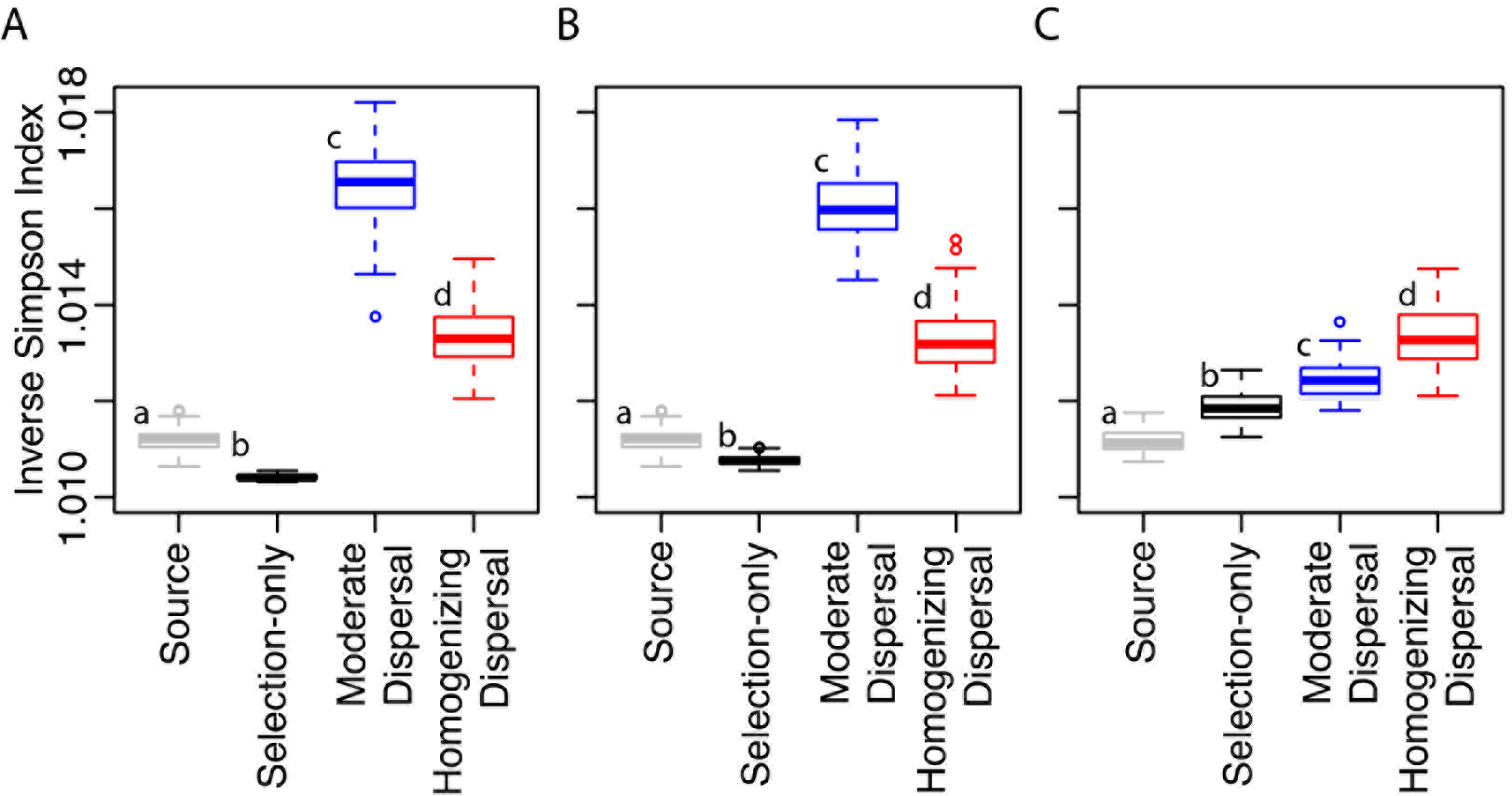
Alpha diversity (inverse Simpson Index) of communities assembled under wide (**A**, niche breadth = 0.175), moderate (**B**, niche breadth = 0.086842105), and narrow (**C**, niche breadth = 0.0075), niches in the mid-point environment (0.476315789). Upper and lower hinges of the box plots represent the 75^th^ and 25^th^ percentiles and whiskers represent 1.5 times the 75^th^ and 25^th^ percentiles, respectively. Colors coincide with labels on the x-axis.

**Figure 3.**
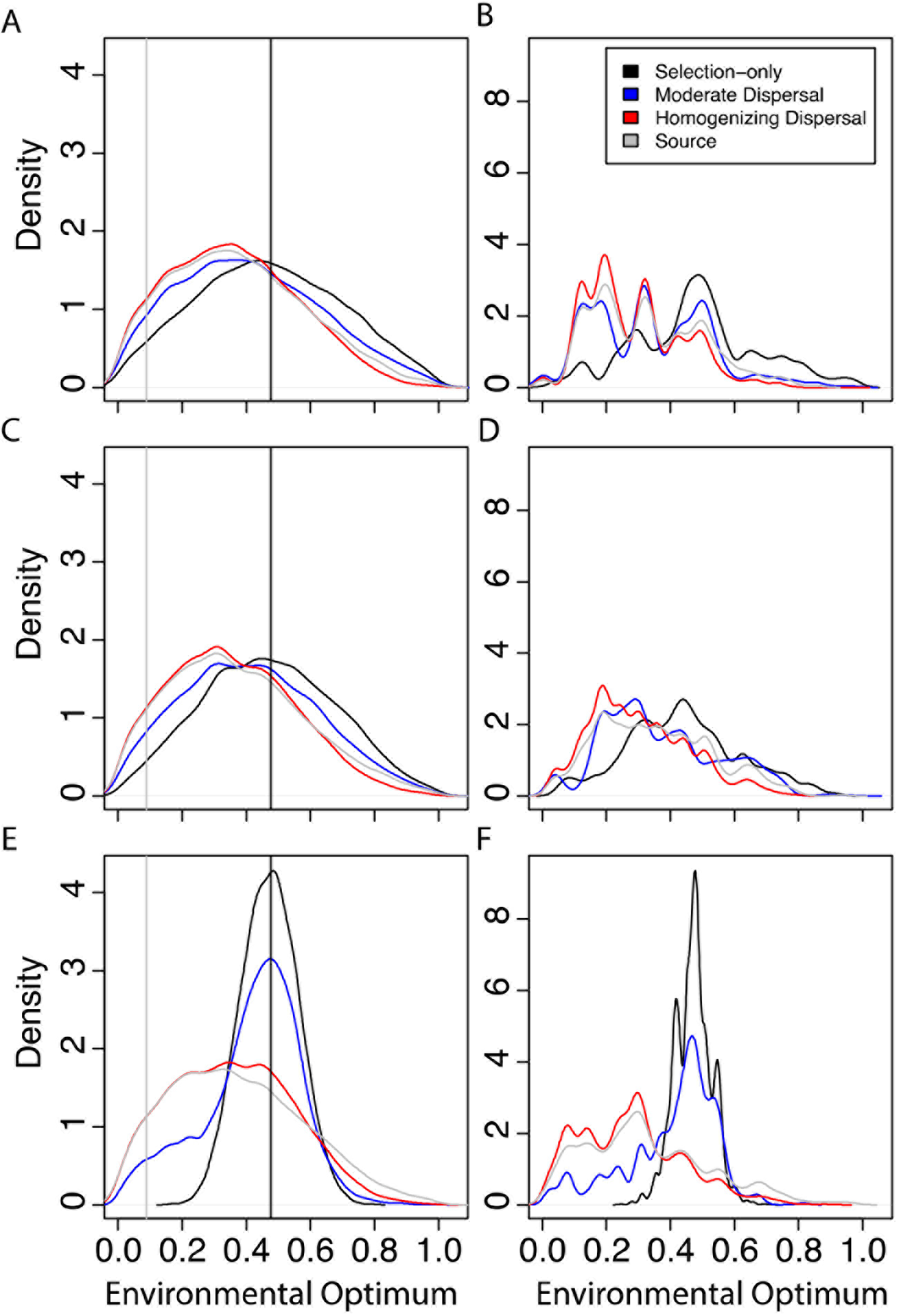
Kernel density of species optima are shown under wide (**A-B**, niche breadth = 0.175), moderate (**C-D**, niche breadth = 0.086842105), and narrow (**E-F**, niche breadth = 0.0075), niches in the mid-point environment (0.476315789, vertical black line). Column 1 displays distributions of species optima without accounting for abundances. Column 2 displays distributions of individuals’ optima. Distributions for the source community and its environment condition (vertical line) are displayed in gray. The same communities were selected as examples to generate Figures 2 and 3.

With the narrowest niche breadths (Figure 2C), we observed a distinct pattern of diversity relative to broader niche breadths (Figure 2A, 2B). Diversity in moderate-dispersal cases decreased substantially as niche breadth narrowed, indicating that moderate levels of dispersal can be overwhelmed when local selective pressures are strong. In contrast, homogenizing dispersal cases maintained consistent levels of diversity across niche breadths and displayed distributions of environmental optima that tracked those of the source community (Figure 3). Diversity in selection-only cases was greatest under the narrowest niche breadth. This was due to only very well-adapted species being part of the community, which led to high abundance across all species in those communities (Figure 3F). For selection-only communities, broader niche breadths resulted in more species with low abundances, and thus lower diversity (cf. black lines in Figure 3B,D,F).

### 3.2 Stochasticity, Microbial Community Composition, and Biogeochemical Function

We found that microbial community assembly history altered the degree to which organisms within a community were adapted to their local environment. Given our assumption of the connection between the degree of adaptation and biogeochemial function (see Methods), assembly history was therefore found to have an indirect effect on biogeochemical function. The environmental optima of taxa in selection-only communities more closely matched their simulated environmental conditions compared to communities assembled with dispersal (Figure 3, 1^st^ column, *p* < 0.001). When the niche breadth was broad (Figure 3A), species’ environmental optima were distributed around the simulated environment under all dispersal cases. However, as niche breadth decreased (Figure 3C, 3E), the species distribution of selection-only cases tightened around the simulated environment, with moderate and homogenizing dispersal cases having a wider distribution. These disparities were maintained when accounting for species abundances (Figure 3B, 3D, 3F), in which selection-only communities had unimodal distributions separate from the source community, while moderate and homogenizing dispersal communities had distributions ranging from unimodal to multi-modal, depending on niche breadth. Dispersal from the source therefore resulted in significant numbers of individuals having large deviations between their environmental optima and the local environmental condition. The large number of maladapted individuals in communities experiencing dispersal from the source resulted in selection-only communities having the highest rates of biogeochemical function, on average, regardless of the simulated environment (Figure 4, *p* < 0.0001).

**Figure 4.**
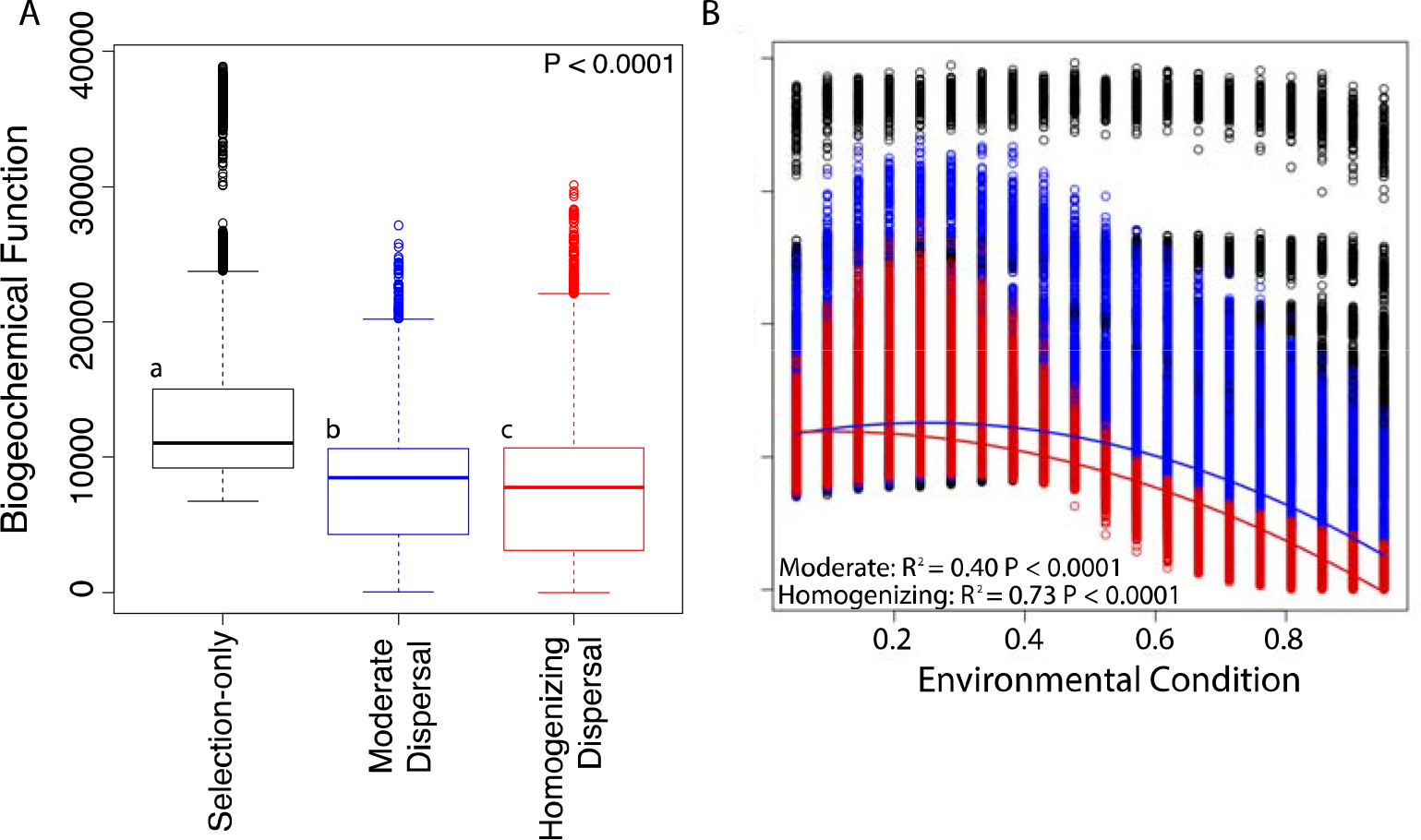
**(A)** Biogeochemical function across dispersal cases. Upper and lower hinges of the box plot represent the 75^th^ and 25^th^ percentiles and whiskers represent 1.5 times the 75^th^ and 25^th^ percentiles, respectively. Different letters indicate statistically significant differences in median values. **(B)** Biogeochemical function across environmental conditions in receiving communities (vertical axis is the same as in panel A). In the selection-only case (black), biogeochemical function did not vary with environmental condition such that no regression line is drawn. With moderate (blue) and homogenizing (red) dispersal, biogeochemical maxima occurred when the receiving community’s environmental condition aligned with the environmental optima of species in the source community (compare to Figure 3). For these two cases, quadratic regression was used and resulting models are shown as solid lines (statistics provided).

In natural systems, microbial community compositional differences can be due to competitive dynamics that select for organisms based on their niche optima (determinism) [46,47] and to immigration of new taxa from the regional species pool (stochasticity) [7,31,48]. Strong local selective pressures can lead to more fit species and enhanced biogeochemistry [7]. Due to the lack of immigrating maladapted species in the selection-only simulations, biogeochemical rates were maintained regardless of the difference between source and receiving community environments. This indicates that biogeochemical function can be enhanced by species adaptation to local conditions. Indeed, a plethora of literature demonstrates that environmental features such as pH [24], nutrients [49], and salinity [25,26] impact microbial community structure and biogeochemical function, and our results indicate that the linkage between community structure and function is due to microbial adaptation to local conditions.

Our results also indicate that when immigrating microorganisms are derived from environments that differ from the receiving community (e.g., dispersal across steep geochemical gradients), biogeochemical function may be suppressed. When we included dispersal from a source community, greater differences between the source and receiving communities led to decreases in biogeochemical function in the receiving communities (Figure 4B, *p* < 0.0001), and this effect became more pronounced as the rate of dispersal increased. Natural systems are influenced by some combination of dispersal and selection and our results indicate that function is maximized when dispersal is minimized and selection is maximized.

Dispersal had the greatest influence on biogeochemical function when niche breadth was narrow (Figure 5). The biogeochemical function of selection-only communities in comparison to homogenizing-dispersal communities was greatest under the narrowest niche breadth (0.0075) and rapidly decreased when transitioning to broader niche breadths. Selection-only communities simulated with narrow niches are comprised of specialist species that can generate high biogeochemical rates and that are well adapted to their local environment. Increasing niche breadth results in the assembly of species with a broader range of environmental optima and that generate lower biogeochemical rates even if their environmental optimum matches the environmental condition (see Methods for a discussion of this assumed trade-off). Thus, high rates of dispersal combined with narrow niche breadths causes replacement of high-functioning specialist organisms with maladapted taxa, thereby significantly reducing community-level biogeochemical function. When niche breadth is broader, immigrating organisms replace lower-functioning organisms (i.e., generalists), resulting in a smaller decreased in community biogeochemical function. We note that our model does not represent dispersal-competition tradeoffs [18], nor does it explicitly represent organismal interactions; exploring the influence of these features would be an interesting extension of the model presented here.

**Figure 5.**
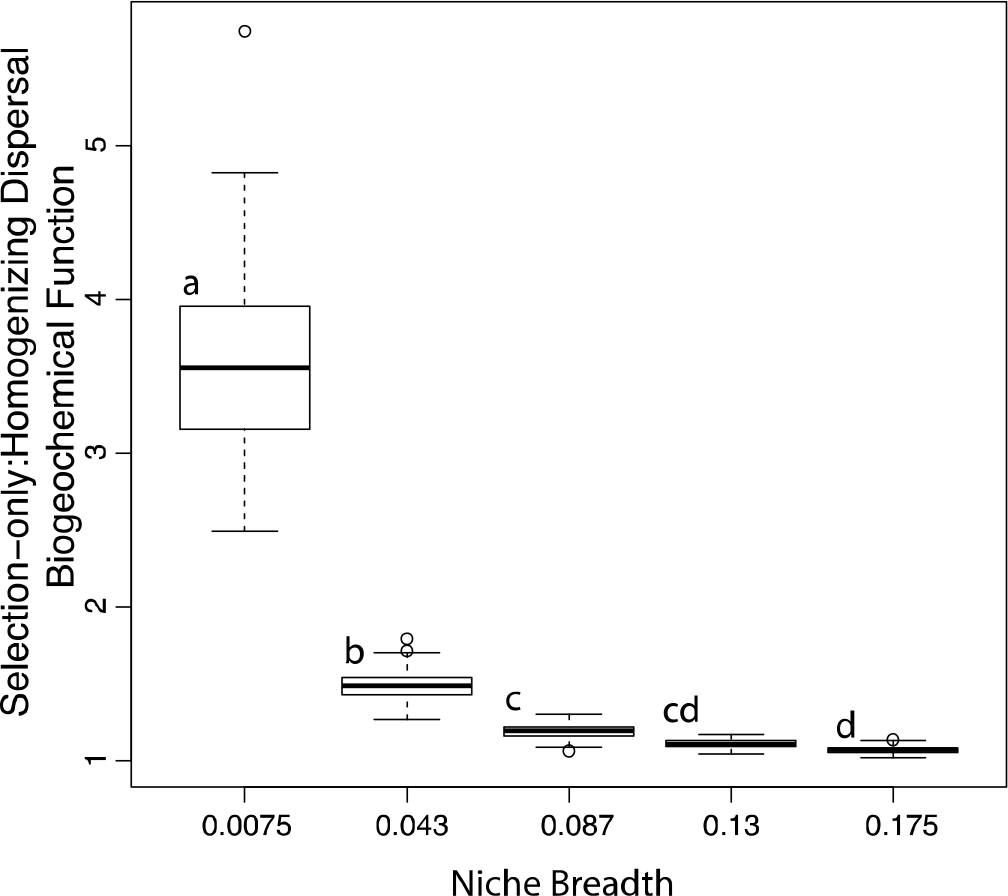
The ratio of biogeochemical function in selection-only cases to homogenizing dispersal cases across 5 niche breadths that span the entire parameter range (0.0075 to 0.175). Each column represents all replicates across all environments for a given niche breadth. Different letters indicate statistically significant differences in mean values. Upper and lower hinges of the box plots represent the 75^th^ and 25^th^ percentiles and whiskers represent 1.5 times the 75^th^ and 25^th^ percentiles, respectively.

Regardless of dispersal, simulations with broader niche breadth led to lower rates of biogeochemical function, supporting a tradeoff between communities comprised of specialist vs. generalist species [50–52], Previous work in microbial systems has posited life-history tradeoffs between specialist vs. generalist species, whereby specialists expend more energy to establish their niches but function at higher levels once established [53]. Specialist species have also been found to be more sensitive to changes in the environment due to strong adaptation to their local environment, with generalists being more resilient to change [54–57]. While we do not address temporal dynamics in our model, the separation of biogeochemical function based on niche breadth indicates a central role for the balance of specialist vs. generalist microorganisms within a community in determining function, regardless of prevailing environmental conditions.

### 3.3 Impact of Assembly Processes on Biogeochemical Function

We also observed that niche breadth within the receiving community was a key parameter in dictating biogeochemical function when environmental conditions (and thus selective pressures) differed between source and receiving communities. In cases without dispersal, biogeochemical function was dictated entirely by niche breadth regardless of differences in selective environments (as inferred from βNTI) between source and receiving communities (Figure 6A, 6D). Selective pressures in the selection-only receiving communities were most dissimilar to the source community (βNTI > 2) in simulations with both narrow niche breadth and environmental conditions that were very different from the source community (Figure 6A). This relationship was also apparent (but weaker) in simulations with an intermediate amount of dispersal (Figure 6B). In receiving communities with high rates of dispersal, stochasticity (|βNTI| < 2) was the dominant process regardless of niche breadth or environmental condition in the receiving community (Figure 6C).

**Figure 6.**
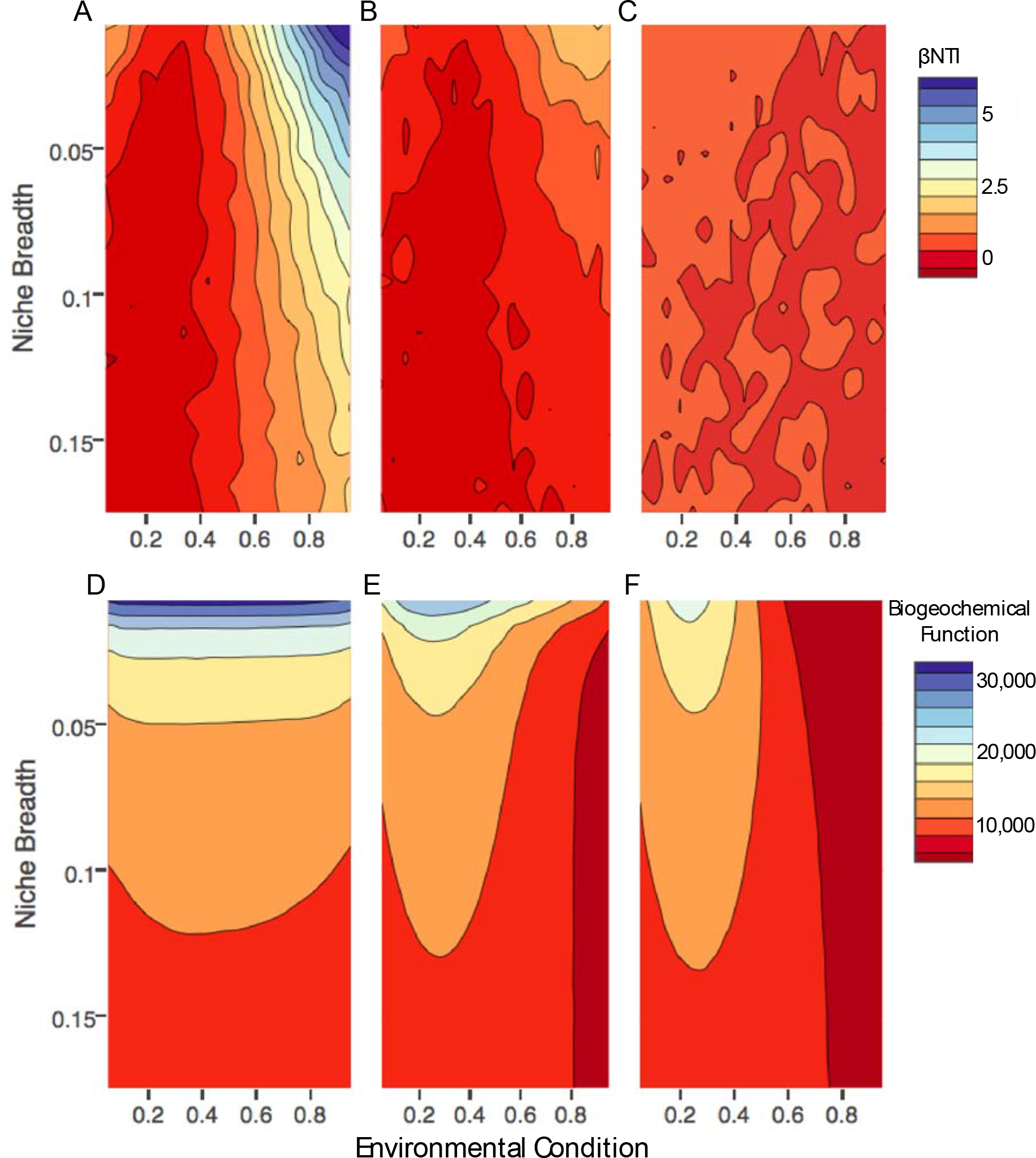
Interpolated contour plots showing average βNTI **(A-C)** and biogeochemical function **(D-F)** for each dispersal case across all parameter combinations, interpolations are based on parameter combinations at each of 20 evenly spaced values across each axis. Values of βNT1 that are further from 0 indicate increasing influences of deterministic assembly (and decreasing stochasticity). **(A)** and **(D)** depict selection-only communities; **(B)** and **(E)** depict moderate dispersal communities; and **(C)** and **(F)** depict homogenizing dispersal communities.

Across the full parameter space defined by niche breadth and environmental condition, cases with moderate and homogenizing dispersal were generally characterized by a dominance of stochasticity (Figure 6B, 6C). This increase in stochasticity relative to selection-only cases corresponded to decreased biogeochemical function. This was particularly true as the environment diverged from the source community (Figure 6D, 6E, 6F). Biogeochemical function in these cases was also negative correlated to niche breadth (*i.e.*, highest under narrow niche breadths), revealing higher functioning of specialist organisms regardless of assembly processes.

Given these apparent associations between assembly processes and biogeochemical function, we directly examined differences in relationships between βNTI and biogeochemical function across a range of environments and niche breadths (Figure 7). Our results suggest that microbial assembly processes may exert the most influence over biogeochemical function when there is significant variation in the relative contributions of deterministic and stochastic processes among communities. We found the strongest relationships between βNTI and function when environmental conditions were dissimilar to the source community, regardless of niche breadth (Figure 7G-I). βNTI had the greatest range in these scenarios, reflecting substantial variation in the contribution of stochastic and deterministic processes. By contrast, scenarios with environments more similar to the source environment had little variation in assembly processes and no relationship between βNTI and biogeochemical function (Figure 7A-F).

**Figure 7.**
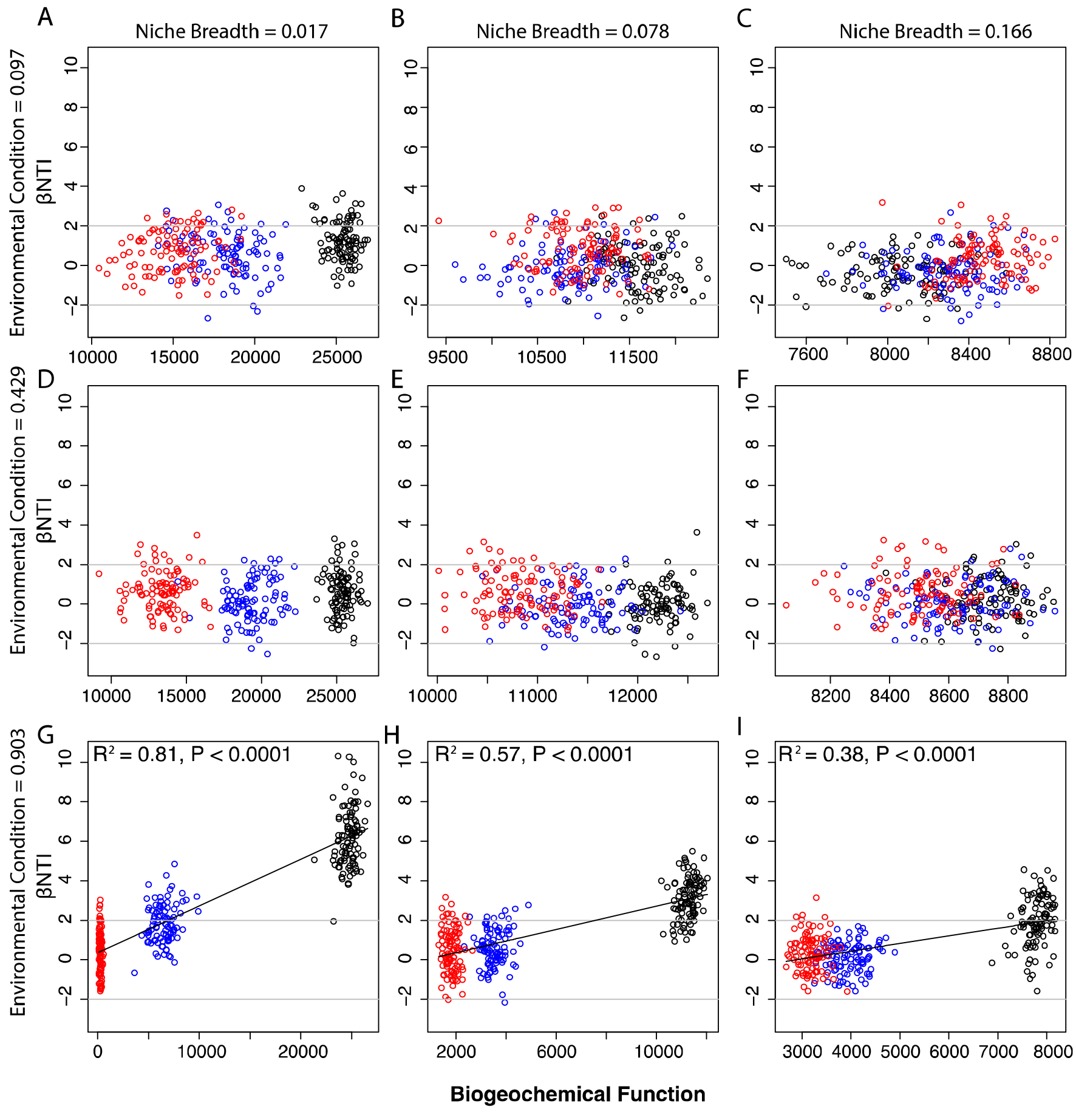
Relationship between βNT1 and biogeochemical function across different niche breadths (columns) and different environmental conditions of the receiving communities (rows). Values of βNT1 that are further from 0 indicate increasing influences of deterministic assembly (and decreasing stochasticity). Horizontal gray lines indicate significance thresholds of −2 and +2. Relationships were evaluated with linear regression; fitted models are shown as black lines and statistics are provided on each panel. Panels without regression models had non-significant (*p* > 0.05) relationships. Note that the vertical axis is scaled the same across panels, but the horizontal axis is not. Black, blue, and red symbols indicate selection-only, moderate dispersal, and homogenizing dispersal scenarios, respectively.

Variation in the balance of stochastic and deterministic assembly processes is prevalent in natural systems [2,7,16,37], as most ecosystems experience spatially and/or temporally variable rates of dispersal. For example, hydrologic connectivity facilitates microbial dispersal and differs with physical matrix structure in soils and sediments. We therefore pose that variation in βNTI may be an effective tool for predicting biogeochemical function when biotic and abiotic conditions lead to a mixture of stochastic and deterministic assembly processes. Natural systems have repeatedly shown such a mixture, and previous field observations have revealed connections between βNTI and biogeochemical function [2,14,58]. These outcomes support our model-based inference that βNTI—as a proxy for assembly processes—offers a practical means to inform models that represent the effects of ecological processes on biogeochemical function.

While our results suggest that maladapted immigrating organisms decrease biogeochemical function, it is important to note that stochasticity may offer buffering capacity that maintains or increases biogeochemical function relative to well-adapted deterministic communities in the context of future environmental perturbations not simulated here [54]. Stochastic spatial processes, such as dispersal, may lead to coexistence of species with different environmental optima resulting in a community that can rapidly adapt to changing environment conditions and maintain biogeochemical function in the face of perturbation. Researchers have long demonstrated positive relationships between biodiversity and ecosystem function in both macrobial [59,60] and microbial [61–63] systems, and new work has highlighted the role of stochasticity in maintaining this connection [64]. Conversely, a lack of stochasticity may result in species so specialized to a given environment that they are vulnerable to environmental change [54]. While these communities would putatively exhibit high rates of biogeochemical function under stable environmental conditions, their function would plummet in response to perturbation, akin to observations of a tradeoff between function and vulnerability in plant communities [20,65].

### 3.4 Implications for Ecosystem Models

The cumulative impacts of ecological processes through time and how they relate to ecosystem-level processes is an emerging research frontier in ecosystem science [2,50,66,67]. We reveal how stochastic community assembly can decrease adaptation to local environments and, in turn, decrease biogeochemical function. Our modelling approach demonstrates plausible outcomes of microbial assembly processes on ecosystem functioning, and integrating this knowledge with factors such as historical abiotic conditions, competitive dynamics, and life-history traits could substantially improve ecosystem model predictions.

Previous work by Hawkes and Keitt [50] laid a theoretical foundation for incorporating time-integrated ecological processes into predictions of biogeochemical function. They demonstrate that community-level microbial functions are the accretion of individual life-histories that determine population growth, composition, and fitness. However, they acknowledge their exclusion of dispersal processes from their models and do not explicitly consider stochasticity in their analysis. Hawkes and Keitt [50] therefore provide a foundation for future research and call for a holistic understanding of historical processes on microbial function, with a particular emphasis on the underlying mechanisms generating these trends. Our work enhances this framework by demonstrating that community assembly processes are integral to knowledge of biogeochemical function in natural systems.

Microbially-explicit models (e.g., MIMICS, MEND) are rapidly becoming more sophisticated and are readily amenable to modules that represent ecological assembly processes [68,69]. As models begin to consider microbial ecology, there is a need to decipher linkages among spatiotemporal microbial processes and ecosystem-level biogeochemical function. We propose that new microbially-explicit models should go beyond microbial mechanisms at a given point in time or space, and building upon the foundation laid by Hawkes and Keitt [50], incorporate ecological dynamics that operate across longer time scales to influence biogeochemical function. Although there are many available avenues to merge modelling efforts in microbial ecology and ecosystem science, there is little debate that integrated models will increase the accuracy of predictions in novel future environments.

### 3.5 Conclusions

We demonstrate the influence of ecological assembly processes on biogeochemical function. Specifically, we show that dispersal can increase the abundance of maladapted organisms in a community, and in turn, decrease biogeochemical function. This impact is strongest when organismal niche breadth is narrow. We also pose that the explanatory power of microbial assembly processes on biogeochemical function is greatest when there is variation in the contributions of stochastic and deterministic processes across a collection of local communities within a broader system of interest. While our work is an encouraging advancement in understanding relationships between ecology and biogeochemistry, a key next step is incorporating assembly processes into emerging model frameworks that explicitly represent microbes and that mechanistically represent biogeochemical reactions.

## Acknowledgments

We thank Andrew Pitman for help editing manuscript text and Nathan Johnson for graphical assistance. This research was supported by the US Department of Energy (DOE), Office of Biological and Environmental Research (BER), as part of Subsurface Biogeochemical Research Program’s Scientific Focus Area (SFA) at the Pacific Northwest National Laboratory (PNNL). PNNL is operated for DOE by Battelle under contract DE-AC06-76RLO 1830. This research was performed using Institutional Computing at PNNL.

## Author Contributions

E.B.G and J.C.S conceived and designed this work; E.B.G. performed the simulations, analyzed the data, and wrote the manuscript; J.C.S. contributed to manuscript revisions and code development.

## Conflicts of Interest

The authors declare no conflicts of interest.

## References

1. Wallenstein, M.D.; Hall, E.K. A trait-based framework for predicting when and where microbial adaptation to climate change will affect ecosystem functioning. Biogeochemistry 2012, 109, 35–47.

2. Graham, E.B.; Crump, A.R.; Resch, C.T.; Fansler, S.; Arntzen, E.; Kennedy, D.W.; Fredrickson, J.K.; Stegen, J.C. Coupling spatiotemporal community assembly processes to changes in microbial metabolism. Frontiers in Microbiology 2016, 7.

3. Graham, E.B.; Knelman, J.E.; Schindlbacher, A.; Siciliano, S.; Breulmann, M.; Yannarell, A.; Beman, J.; Abell, G.; Philippot, L.; Prosser, J. Microbes as engines of ecosystem function: When does community structure enhance predictions of ecosystem processes? Frontiers in microbiology 2016, 7.

4. Martiny, J.B.; Jones, S.E.; Lennon, J.T.; Martiny, A.C. Microbiomes in light of traits: A phylogenetic perspective. Science 2015, 350, aac9323.

5. Bier, R.L.; Bernhardt, E.S.; Boot, C.M.; Graham, E.B.; Hall, E.K.; Lennon, J.T.; Nemergut, D.R.; Osborne, B.B.; Ruiz-González, C.; Schimel, J.P. Linking microbial community structure and microbial processes: An empirical and conceptual overview. FEMS microbiology ecology 2015, 91.

6. Rocca, J.D.; Hall, E.K.; Lennon, J.T.; Evans, S.E.; Waldrop, M.P.; Cotner, J.B.; Nemergut, D.R.; Graham, E.B.; Wallenstein, M.D. Relationships between protein-encoding gene abundance and corresponding process are commonly assumed yet rarely observed. The ISME journal 2015, 9, 1693.

7. Nemergut, D.R.; Schmidt, S.K.; Fukami, T.; O’Neill, S.P.; Bilinski, T.M.; Stanish, L.F.; Knelman, J.E.; Darcy, J.L.; Lynch, R.C.; Wickey, P. Patterns and processes of microbial community assembly. Microbiology and Molecular Biology Reviews 2013, 77, 342–356.

8. Nemergut, D.R.; Shade, A.; Violle, C. When, where and how does microbial community composition matter? Frontiers in microbiology 2014, 5.

9. Pholchan, M.K.; Baptista, J.d.C.; Davenport, R.J.; Sloan, W.T.; Curtis, T.P. Microbial community assembly, theory and rare functions. Frontiers in microbiology 2013, 4.

10. Cardinale, B.J.; Wright, J.P.; Cadotte, M.W.; Carroll, I.T.; Hector, A.; Srivastava, D.S.; Loreau, M.; Weis, J.J. Impacts of plant diversity on biomass production increase through time because of species complementarity. Proceedings of the National Academy of Sciences 2007, 104, 18123–18128.

11. Cadotte, M.W.; Carscadden, K.; Mirotchnick, N. Beyond species: Functional diversity and the maintenance of ecological processes and services. Journal of applied ecology 2011, 48,1079–1087.

12. Cadotte, M.W. Dispersal and species diversity: A meta-analysis. The American Naturalist 2006, 167, 913–924.

13. Cardinale, B.J. Biodiversity improves water quality through niche partitioning. Nature 2011, 472, 86.

14. Graham, E.B.; Crump, A.R.; Resch, C.T.; Fansler, S.; Arntzen, E.; Kennedy, D.W.; Fredrickson, J.K.; Stegen, J.C. Deterministic influences exceed dispersal effects on hydrologically-connected microbiomes. Environmental Microbiology 2017, 19, 1552–1567.

15. Stegen, J.C.; Lin, X.; Fredrickson, J.K.; Konopka, A.E. Estimating and mapping ecological processes influencing microbial community assembly. Frontiers in microbiology 2015, 6.

16. Vellend, M. Conceptual synthesis in community ecology. The Quarterly review of biology 2010, 85, 183–206.

17. Leibold, M.A.; Holyoak, M.; Mouquet, N.; Amarasekare, P.; Chase, J.M.; Hoopes, M.F.; Holt, R.D.; Shurin, J.B.; Law, R.; Tilman, D. The metacommunity concept: A framework for multi-scale community ecology. Ecology letters 2004, 7, 601–613.

18. Tilman, D.; Lehman, C.L.; Bristow, C.E. Diversity-stability relationships: Statistical inevitability or ecological consequence? The American Naturalist 1998, 151, 277–282.

19. Doak, D.F.; Bigger, D.; Harding, E.; Marvier, M.; O’malley, R.; Thomson, D. The statistical inevitability of stability-diversity relationships in community ecology. The American Naturalist 1998, 151, 264–276.

20. Tilman, D. The ecological consequences of changes in biodiversity: A search for general principles. Ecology 1999, 80, 1455–1474.

21. Strickland, M.S.; Lauber, C.; Fierer, N.; Bradford, M.A. Testing the functional significance of microbial community composition. Ecology 2009, 90, 441–451.

22. Pulliam, H.R. Sources, sinks, and population regulation. The American Naturalist 1988, 132, 652–661.

23. Shmida, A.; Wilson, M. V. Biological determinants of species diversity. Journal of biogeography 1985, 1–20.

24. Fierer, N.; Jackson, R.B. The diversity and biogeography of soil bacterial communities. Proceedings of the National Academy of Sciences of the United States of America 2006, 103, 626–631.

25. Hollister, E.B.; Engledow, A.S.; Hammett, A.J.M.; Provin, T.L.; Wilkinson, H.H.; Gentry, T.J. Shifts in microbial community structure along an ecological gradient of hypersaline soils and sediments. The ISME journal 2010, 4, 829.

26. Casamayor, E.O.; Massana, R.; Benlloch, S.; Øvreås, L.; Díez, B.; Goddard, V.J.; Gasol, J.M.; Joint, I.; Rodríguez-Valera, F.; Pedrós-Alió, C. Changes in archaeal, bacterial and eukaryal assemblages along a salinity gradient by comparison of genetic fingerprinting methods in a multipond solar saltern. Environmental Microbiology 2002, 4, 338–348.

27. Gotelli, N.J.; Graves, G.R. Null models in ecology. 1996.

28. Cadotte, M.W.; Tucker, C.M. Should environmental filtering be abandoned? Trends in Ecology & Evolution 2017.

29. Hurlbert, A.H.; Stegen, J.C. When should species richness be energy limited, and how would we know? Ecology letters 2014, 17,401–413.

30. Stegen, J.C.; Lin, X.; Fredrickson, J.K.; Chen, X.; Kennedy, D.W.; Murray, C.J.; Rockhold, M.L.; Konopka, A. Quantifying community assembly processes and identifying features that impose them. The ISME journal 2013, 7, 2069.

31. Stegen, J.C.; Lin, X.; Konopka, A.E.; Fredrickson, J.K. Stochastic and deterministic assembly processes in subsurface microbial communities. The ISME journal 2012, 6, 1653.

32. Stegen, J.C.; Hurlbert, A.H. Inferring ecological processes from taxonomic, phylogenetic and functional trait β-diversity. PloS one 2011, 6, e20906.

33. Chase, J.M.; Leibold, M.A. Ecological niches: linking classical and contemporary approaches. University of Chicago Press: 2003.

34. Webb, C.O.; Ackerly, D.D.; Kembel, S.W. Phylocom: Software for the analysis of phylogenetic community structure and trait evolution. Bioinformatics 2008, 24, 2098–2100.

35. Fine, P.V.; Kembel, S.W. Phylogenetic community structure and phylogenetic turnover across space and edaphic gradients in western amazonian tree communities. Ecography 2011, 34, 552–565.

36. Dini-Andreote, F.; Stegen, J.C.; van Elsas, J.D.; Salles, J.F. Disentangling mechanisms that mediate the balance between stochastic and deterministic processes in microbial succession. Proceedings of the National Academy of Sciences 2015, 112, E1326–E1332.

37. Wang, J.; Shen, J.; Wu, Y.; Tu, C.; Soininen, J.; Stegen, J.C.; He, J.; Liu, X.; Zhang, L.; Zhang, E. Phylogenetic beta diversity in bacterial assemblages across ecosystems: Deterministic versus stochastic processes. The ISME journal 2013, 7, 1310.

38. Jurburg, S.D.; Nunes, I.; Stegen, J.C.; Le Roux, X.; Priemé, A.; Sørensen, S.J.; Salles, J.F. Autogenic succession and deterministic recovery following disturbance in soil bacterial communities. Scientific Reports 2017, 7.

39. Veach, A.M.; Stegen, J.C.; Brown, S.P.; Dodds, W.K.; Jumpponen, A. Spatial and successional dynamics of microbial biofilm communities in a grassland stream ecosystem. Molecular ecology 2016, 25,4674–4688.

40. Hill, M.O. Diversity and evenness: A unifying notation and its consequences. Ecology 1973, 54, 427–432.

41. Oksanen, J.; Blanchet, F.G.; Kindt, R.; Legendre, P.; Minchin, P.R.; O’hara, R.; Simpson, G.L.; Solymos, P.; Stevens, M.H.H.; Wagner, H. Package ‘vegan’. Community ecology package, version 2013, 2.

42. Buchkowski, R.W.; Bradford, M.A.; Grandy, A.S.; Schmitz, O.J.; Wieder, W.R. Applying population and community ecology theory to advance understanding of belowground biogeochemistry. Ecology letters 2017, 20, 231–245.

43. Wang, G.; Jagadamma, S.; Mayes, M.A.; Schadt, C.W.; Steinweg, J.M.; Gu, L.; Post, W.M. Microbial dormancy improves development and experimental validation of ecosystem model. The ISME journal 2015, 9, 226.

44. Wieder, W.; Grandy, A.; Kallenbach, C.; Taylor, P.; Bonan, G. Representing life in the earth system with soil microbial functional traits in the mimics model. Geoscientific Model Development 2015, 8, 1789–1808.

45. Connell, J.H. Diversity in tropical rain forests and coral reefs. Science 1978, 199,1302–1310.

46. DeAngelis, K.M.; Silver, W.L.; Thompson, A.W.; Firestone, M.K. Microbial communities acclimate to recurring changes in soil redox potential status. Environmental Microbiology 2010, 12, 3137–3149.

47. Cregger, M.A.; Schadt, C.W.; McDowell, N.G.; Pockman, W.T.; Classen, A.T. Response of the soil microbial community to changes in precipitation in a semiarid ecosystem. Applied and environmental Microbiology 2012, 78, 8587–8594.

48. Green, J.; Bohannan, B.J. Spatial scaling of microbial biodiversity. Trends in ecology & evolution 2006, 21, 501–507.

49. Schimel, D.S.; Braswell, B.; McKeown, R.; Ojima, D.S.; Parton, W.; Pulliam, W. Climate and nitrogen controls on the geography and timescales of terrestrial biogeochemical cycling. Global Biogeochemical Cycles 1996, 10, 677–692.

50. Hawkes, C.V.; Keitt, T.H. Resilience vs. Historical contingency in microbial responses to environmental change. Ecology letters 2015, 18, 612–625.

51. Wilson, D.S.; Yoshimura, J. On the coexistence of specialists and generalists. The American Naturalist 1994, 144, 692–707.

52. Kassen, R. The experimental evolution of specialists, generalists, and the maintenance of diversity. Journal of evolutionary biology 2002, 15, 173–190.

53. Lennon, J.T.; Aanderud, Z.T.; Lehmkuhl, B.; Schoolmaster, D.R. Mapping the niche space of soil microorganisms using taxonomy and traits. Ecology 2012, 93, 1867–1879.

54. Shade, A.; Peter, H.; Allison, S.D.; Baho, D.L.; Berga, M.; Bürgmann, H.; Huber, D.H.; Langenheder, S.; Lennon, J.T.; Martiny, J.B. Fundamentals of microbial community resistance and resilience. Frontiers in microbiology 2012, 3.

55. Mou, X.; Sun, S.; Edwards, R.A.; Hodson, R.E.; Moran, M.A. Bacterial carbon processing by generalist species in the coastal ocean. Nature 2008, 451, 708.

56. Langenheder, S.; Lindström, E.S.; Tranvik, L.J. Weak coupling between community composition and functioning of aquatic bacteria. Limnology and Oceanography 2005, 50, 957–967.

57. Allison, S.D.; Martiny, J.B. Resistance, resilience, and redundancy in microbial communities. Proceedings of the National Academy of Sciences 2008, 105, 11512–11519.

58. Stegen, J.C.; Fredrickson, J.K.; Wilkins, M.J.; Konopka, A.E.; Nelson, W.C.; Arntzen, E.V.; Chrisler, W.B.; Chu, R.K.; Danczak, R.E.; Fansler, S.J. Groundwater-surface water mixing shifts ecological assembly processes and stimulates organic carbon turnover. Nature communications 2016, 7.

59. Cardinale, B.J.; Matulich, K.L.; Hooper, D.U.; Byrnes, J.E.; Duffy, E.; Gamfeldt, L.; Balvanera, P.; O’Connor, M.I.; Gonzalez, A. The functional role of producer diversity in ecosystems. American journal of botany 2011, 98, 572–592.

60. Hooper, D.U.; Adair, E.C.; Cardinale, B.J.; Byrnes, J.E.; Hungate, B.A.; Matulich, K.L.; Gonzalez, A.; Duffy, J.E.; Gamfeldt, L.; O’Connor, M.I. A global synthesis reveals biodiversity loss as a major driver of ecosystem change. Nature 2012, 486, 105–108.

61. Bell, T.; Newman, J.A.; Silverman, B.W.; Turner, S.L.; Lilley, A.K. The contribution of species richness and composition to bacterial services. Nature 2005, 436, 1157.

62. Langenheder, S.; Bulling, M.T.; Solan, M.; Prosser, J.I. Bacterial biodiversity-ecosystem functioning relations are modified by environmental complexity. PloS one 2010, 5, e10834.

63. Levine, U.Y.; Teal, T.K.; Robertson, G.P.; Schmidt, T.M. Agriculture’s impact on microbial diversity and associated fluxes of carbon dioxide and methane. The ISME journal 2011, 5, 1683.

64. Knelman, J.E.; Nemergut, D.R. Changes in community assembly may shift the relationship between biodiversity and ecosystem function. Frontiers in microbiology 2014, 5.

65. Tilman, D.; Reich, P.B.; Knops, J.; Wedin, D.; Mielke, T.; Lehman, C. Diversity and productivity in a longterm grassland experiment. Science 2001, 294, 843–845.

66. Evans, S.E.; Wallenstein, M.D. Soil microbial community response to drying and rewetting stress: Does historical precipitation regime matter? Biogeochemistry 2012, 109, 101–116.

67. Fukami, T. Historical contingency in community assembly: Integrating niches, species pools, and priority effects. Annual Revieiv of Ecology, Evolution, and Systematics 2015, 46.

68. Wang, G.; Post, W.M.; Mayes, M.A. Development of microbial-enzyme-mediated decomposition model parameters through steady-state and dynamic analyses. Ecological Applications 2013, 23, 255–272.

69. Wieder, W.; Grandy, A.; Kallenbach, C.; Bonan, G. Integrating microbial physiology and physio-chemical principles in soils with the microbial-mineral carbon stabilization (mimics) model. Biogeosciences 2014, 11, 3899–3917.

